# pysigscore: gene signatures scoring across bulk and single-cell transcriptomics

**DOI:** 10.64898/2026.08.04.742537

**Authors:** Tommaso Giacomello, Saveria Mazzara, Gennaro Abbruzzese, Alessandro Barberis, Andrea Tangherloni, Francesca M Buffa

## Abstract

**Summary:** High-throughput transcriptomics has made gene signatures central to interpreting gene expression data, with applications in diagnosis, prognosis, and prediction. Quantifying signature activity and assessing its robustness remain challenging because scoring methods primarily rely on various assumptions, and no single approach is universally optimal. Here, we present `pysigscore`, a Python framework for gene set scoring in bulk and single-cell RNA-seq data. `pysigscore` integrates 18 built-in scoring methods with a fully customisable scorer, allowing users to define and benchmark new scoring functions. It also provides reliability analyses, including p-value estimation and leave-one-out experiments, to assess the significance of scores and gene-level contributions. We validated `pysigscore` on the CCLE, TCGA, and PBMC datasets, recovering the expected enrichment in liver, hypoxia, inflammatory, and cell-cycle signatures.

**Availability and Implementation:** Source code is available at https://github.com/bioinformatics-hub/pysigscore. Contact

**Supplementary information:** Supplementary data are available at Bioinformatics online.

## Introduction

Advances in transcriptomics have generated large-scale expression datasets, from bulk RNA-seq cohorts to single-cell atlases, enabling the widespread use of gene signatures to study biological function and making them a central tool in biomedical research. Gene-set analysis is now routinely applied to quantify signature activity in each sample using a numerical score, enabling the characterisation of tumour biology, the identification of cell states, and the comparison of molecular programmes across diverse experimental settings (Liberzon et al., 2015). Several scoring strategies have been proposed, including simple summaries of gene-set expression, Z-statistics (Kim and Volsky, 2005), and rank-based methods such as ssGSEA (Barbie et al., 2009), GSVA (Hänzelmann et al., 2013), and AUCell (Aibar et al., 2017). These methods differ in their assumptions, robustness, and dependence on background distributions, so no single approach is optimal for all datasets or biological questions. This highlights the need for a framework to compute and compare multiple scoring methods.

`pysigscore` builds on our previous work on signature quality control and scoring, including sigQC for systematic evaluation of individual molecular signatures (Dhawan et al., 2019), and the R package `sigscores` for statistical analysis of multiple signature scores (Barberis and Buffa, 2026). Building on the functionality of `sigscores, pysigscore` extends this framework to Python while introducing several important new capabilities: (*i*) it includes new approaches developed specifically for single-cell transcriptomics, increasing the number of built-in scoring methods to 18; (*ii*) it provides a flexible built-in custom scorer, enabling users to implement and directly compare their own scoring methods with existing approaches within a single unified library; (*iii*) it allows the contribution of individual genes to signature activity to be evaluated; (*iv*) it is optimised for scalability to large transcriptomic datasets; and (*v*) it is directly compatible with the major RNA-seq analysis frameworks, facilitating seamless integration into existing analysis pipelines. Together, these features allow users not only to compute scores but also to compare and assess the robustness of different methods within a single, reproducible framework.

## System and methods

`pysigscore` framework is coordinated by a `Scorer()` object. Starting from user-provided input data, it manages preprocessing, score computation, reliability assessment, and plotting within a unified workflow (see Fig. 1). The workflow requires two main inputs: an expression matrix, from either bulk or scRNA-seq data, and a collection of gene sets. Expression data can be provided as dataframes or as `AnnData` objects (Virshup et al., 2024), compatible with usual scRNA-seq workflows. Gene sets can be supplied as dictionaries or binary matrices, derived from known databases or custom-built.

**Figure 1.**
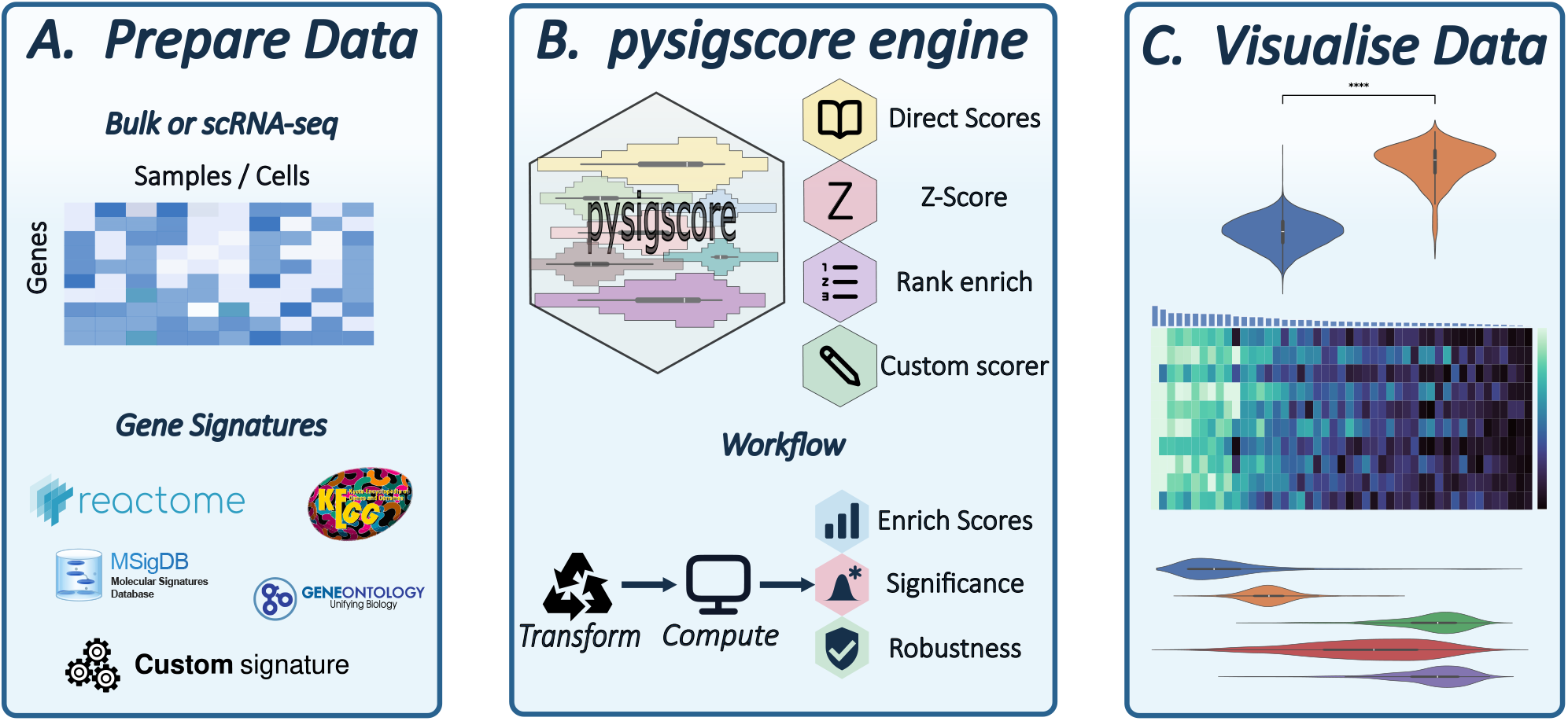
Schematic overview of the `pysigscore` pipeline. (a) It accepts bulk or single-cell RNA-seq data as input, together with a collection of gene signatures derived from established databases or custom-defined by the user. (b) It enables the computation of different scores, along with a significance and robustness assessment. (c) It facilitates the comparison of different scores across samples through dedicated visualisations.

The main entry point of `pysigscore` is the function `compute_scores()`, which applies preprocessing steps to the expression matrix, computes the selected gene-set scores, and optionally evaluates statistical significance and robustness. Data can be considered raw and/or transformed (once or sequentially) to include log transformation, standardisation, min-max scaling, *k*-Nearest-Neighbours (*k*-NN) smoothing (Kamimoto et al., 2023), as well as step and quantile transformations (Barberis and Buffa, 2026). For each selected score and transformation combination, `compute _scores()` returns a score matrix with samples as rows and gene sets as columns. Thus, each entry represents the activity of a gene set in a given sample, computed from the sample’s expression profile and the genes in that signature. Users may apply post-enrichment transformations (e.g., scaling) to improve comparability across samples or gene sets. When requested, `pysigscore` also generates, for each entry, significance and robustness matrices. Finally, scores, gene sets, and samples can also be visualised and compared with plotting functions.

### Enrichment scores

The current implementation provides 18 built-in scoring methods, which operate on one sample and one gene set at a time, organised into three families: direct, z-score, and rank-based methods (Table 1). Fourteen direct methods compute the score only from the subset of values corresponding to the signature genes, providing fast and interpretable measures. In contrast, Z-score uses the full expression profile of the same sample as background, comparing the mean expression of the gene set with the sample-wise distribution. Finally, AUCell, ssGSEA, and GSVA are rank-based methods that assess whether signature genes are enriched among the highest-ranked genes within the same sample.

**Table 1.**
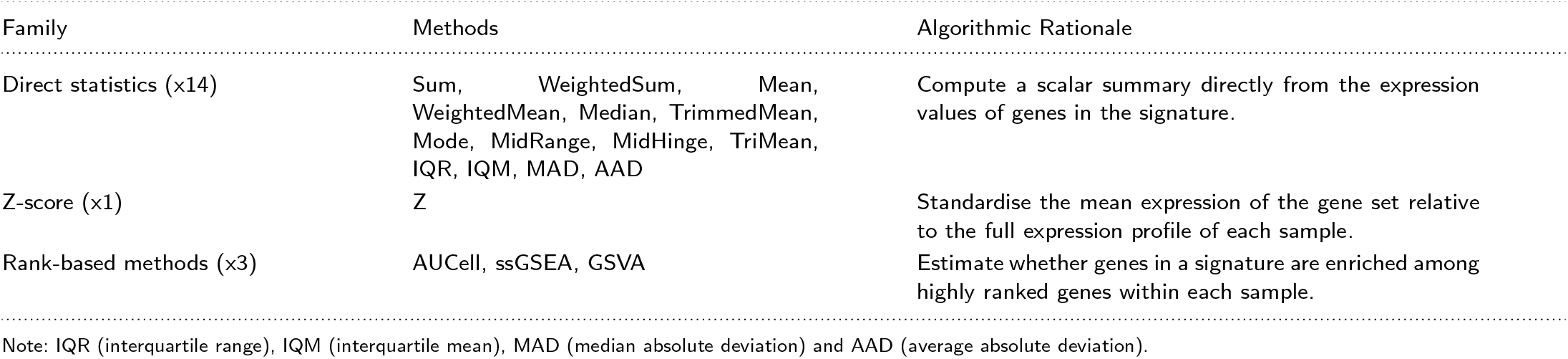
Overview of scoring families, implemented methods, and algorithmic rationale in pysigscore.

### Custom Scorer

The custom scorer allows users to define new custom scoring functions without modifying the `pysigscore` source code. A custom Python function must take exactly two arguments: `arr_geneset`, the expression vector of the current gene set, and `arr_full`, the full expression vector of the same sample or cell.

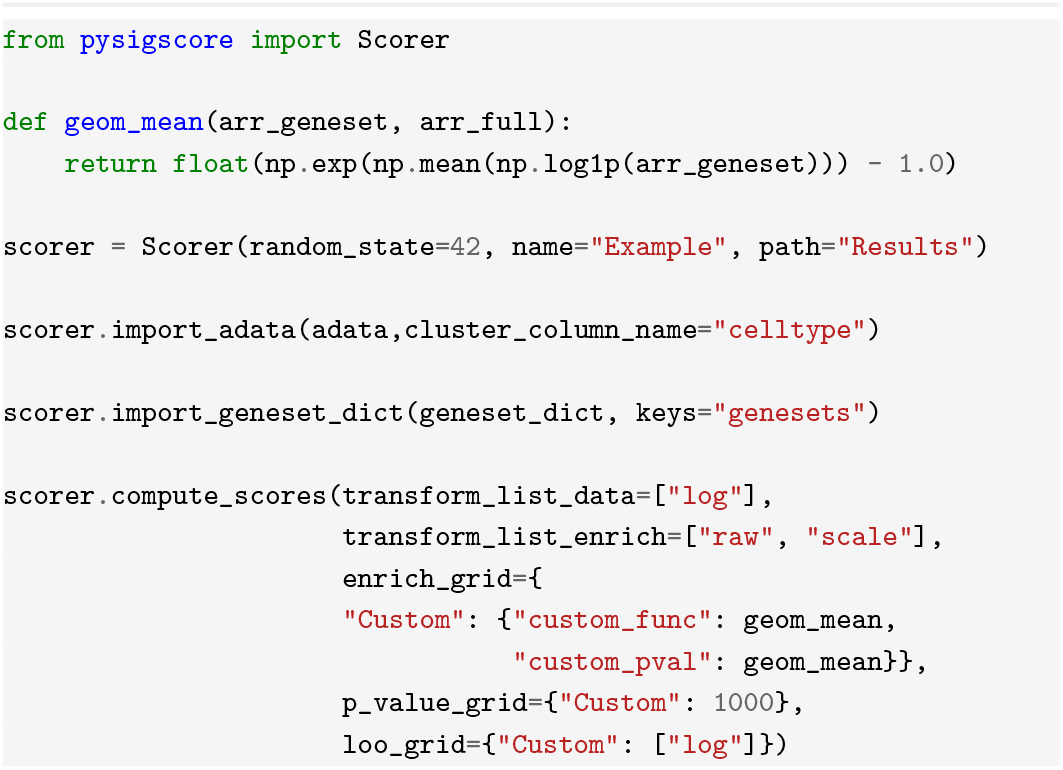

### P-value computation

`pysigscore` assesses the significance of each entry in the score matrix through the `p_value_grid` argument. For direct methods, this argument specifies the number of random, size-matched gene sets sampled per sample and signature to generate a null distribution; nominal p-values then estimate how often null scores exceed the observed score, using the following plus-one correction:

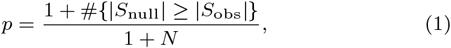

where *S*_obs_ is the observed score, *S*_null_ the null scores, and *N* the number of random signatures). Absolute values account for both directions: a positive (activating) score must exceed the null, a negative (repressing) score must fall below it.

The Z-score method uses a different procedure, as its significance is computed analytically rather than by permutation. Specifically, a two-sided p-value is obtained from the standard normal distribution using the absolute value of the computed Z-score.

For rank-based methods, AUCell and GSVA follow the empirical procedure in Equation 1. While for ssGSEA, p-values are returned by the backend implementation and follow the same general principle, but without the plus-one correction.

### Leave-one-out experiments

The robustness is assessed through leave-one-out (LOO) experiments. For each entry, the score is recomputed by removing each signature gene in turn, and the resulting deviation from the full gene-set score is quantified using mean absolute error (MAE), mean squared error (MSE), or mean absolute percentage error (MAPE). Large deviations may indicate dependence on a few hub genes, whereas small differences support a more evenly weighted signature.

LOO deviations depend on both the scoring method and preprocessing. Direct scores might be more sensitive to highly expressed genes, whereas rank-based scores might reflect overall signature compactness. Raw data emphasise expression magnitude, while standardised data highlight relative gene importance (see Supplementary Material).

For each gene set, `pysigscore` returns a sample-by-gene LOO matrix highlighting each gene deviation. Then, merging all the signature results yields the usual sample-by-gene-set matrix.

### Comparison with existing methods

As reported in Table 2, existing Python tools address specific aspects of gene-set scoring, but none provides a comprehensive framework like `pysigscore`. For instance, `gseapy` implements methods such as ssGSEA and GSVA (Fang et al., 2023), and `pysigscore` uses it as the backend for these approaches. `Scanpy` provides `sc.tl.score_genes()` (Wolf et al., 2018), a corrected mean score integrated with scRNA-seq workflows. `decoupler` (Badia-I-Mompel et al., 2022) offers multiple network and model-based methods for regulator and pathway activity inference. In contrast, `pysigscore` provides a broader end-to-end framework by integrating multiple scoring families, enabling direct comparison across scores, together with a user-defined custom scorer, reliability analyses, and plotting functions within a single Python workflow.

**Table 2.** Comparison of pysigscore with the existing Python packages.

| Feature | `pysigscore` | `gseapy` | `decoupler` | `sc.tl.score_genes()` |
| --- | --- | --- | --- | --- |
| Scoring methods | 18 | ssGSEA, GSVA | 11 | 1 |
| Custom scorer | Yes | No | No | No |
| Method classes | Summary, Z-score, rank-based | Rank-based | Network/activity inference | Mean-based |
| P-value support | Yes | Method-dependent | Method-dependent | No |
| Leave-one-out analysis | Yes | No | No | No |
| Integrated plotting/export | Yes | Yes | Yes | No |

## Application

We validated pysigscore on both bulk RNA-seq and scRNA-seq datasets. Bulk analyses used CCLE cancer cell lines (Ghandi et al., 2019) and TCGA primary tumours (Weinstein et al., 2013), whereas single-cell analyses used PBMC CITE-seq data (Stuart et al., 2019). We evaluated whether Hallmark gene sets (Liberzon et al., 2015) captured the expected tissue-specific, tumour-associated, and immune cell-state signals.

In bulk cancer data, `pysigscore` recovered the liver enrichment of `HALLMARK_BILE_ACID_METABOLISM`. In CCLE, the signal was consistent across scoring methods and liver cell lines showed higher activity than non-liver cell lines (see Figs S1–S4). The same biological pattern was observed in TCGA, where liver tumours were enriched relative to non-liver tumours, with score significance and LOO analyses further supporting the results (Figs S5–S9).

As a second validation example, we compared two independent signatures representing the same hypoxic state (i.e., `HALLMARK_HYPOXIA` and `BUFFA_HYPOXIA_METAGENE` (Buffa et al., 2010)). In CCLE, the two signatures showed concordant behaviour across representative scoring methods, indicating that they captured the same direction of hypoxia-associated variation (see Figs S10–S13). This agreement was reproduced in TCGA, where the two signatures again showed concordant activity across tumour samples (see Figs S14–S17).

In single-cell PBMC data, the `HALLMARK_INFLAMMATORY_RESPONSE` was enriched in *CD14* ^+^ monocytes but not in naive B cells, confirming the inflammatory role of monocytes. Multiple scores were correlated; therefore, the claim was observed across representative methods and was supported by p-values, statistical comparisons, and LOO analyses (see Figs S18–S22). As a final validation in PBMCs, the `HALLMARK_G2M_CHECKPOINT` enrichment was significantly higher in proliferating *CD4* and *CD8* T cells than in their respective naive counterparts. Despite some score-dependent differences, representative methods consistently separated proliferating from naive populations, with statistical comparison and LOO analysis supporting the cell-cycle signal (see Figs S22–S26).

## Discussion and Conclusions

`pysigscore` fills a practical gap in transcriptomics workflows by integrating multiple scoring methods, statistical interpretation, robustness diagnostics, user-defined methods, and visualisation within a single Python framework. Whereas existing tools typically focus on individual scores or workflows tailored to specific data types, `pysigscore` enables users to compare alternative definitions of gene set activity and assess whether biological conclusions are consistent across methods. This breadth of analysis necessarily introduces a computational cost. Nevertheless, under the evaluated configurations, including 100 permutations and LOO analysis, `pysigscore` completed the CCLE and TCGA analyses in 3 h 40 min and 1 d 1 h 20 min, with peak memory requirements of 4.07 and 25.61 GB, respectively. The PBMC analysis required 3 d 10 h and 83.24 GB of RAM on a single CPU core, reflecting the increased demands of large single-cell datasets (see Supplementary Material). All the analyses were run using a single CPU core on an Intel Xeon 6517P processor.

Future work will focus on improving computational scalability and extending the framework to other omics data types. Although not explicitly evaluated here, the `AnnData`-based input format is compatible with spatial transcriptomics data, which will be addressed in future validation studies.

## Data and Code availability

Dataset information, code and experimental details are provided in `Supplementary_Information` and `Supplementary_Tables`. Code is available at https://github.com/bioinformatics-hub/pysigscore.

## Author contributions statement

T.G. wrote and maintained the package, performed the analyses, and wrote the initial manuscript draft. S.M. assisted in manuscript revision and figure preparation. G.A., A.B., A.T. and F.M.B. contributed to software design and manuscript revision. F.M.B. conceived and supervised the project.

